# How to improve parameter estimates in GLM-based fMRI data analysis: cross-validated Bayesian model averaging

**DOI:** 10.1101/095778

**Authors:** Joram Soch, Achim Pascal Meyer, John-Dylan Haynes, Carsten Allefeld

## Abstract

In functional magnetic resonance imaging (fMRI), model quality of general linear models (GLMs) for first-level analysis is rarely assessed. In recent work (Soch et al., 2016: “How to avoid mismodelling in GLM-based fMRI data analysis: cross-validated Bayesian model selection”, NeuroImage, vol. 141, pp. 469-489; DOI: 10.1016/j. neuroimage.2016.07.047), we have introduced cross-validated Bayesian model selection (cvBMS) to infer the best model for a group of subjects and use it to guide second-level analysis. While this is the optimal approach given that the same GLM has to be used for all subjects, there is a much more efficient procedure when model selection only addresses nuisance variables and regressors of interest are included in all candidate models. In this work, we propose cross-validated Bayesian model averaging (cvBMA) to improve parameter estimates for these regressors of interest by combining information from all models using their posterior probabilities. This is particularly useful as different models can lead to different conclusions regarding experimental effects and the most complex model is not necessarily the best choice. We find that cvBMS can prevent not detecting established effects and that cvBMA can be more sensitive to experimental effects than just using even the best model in each subject or the model which is best in a group of subjects.

## 1 Introduction

In *functional magnetic resonance imaging* (fMRI), data are most commonly analyzed using *general linear models* (GLMs) which construct a relation between psychologically defined conditions and the measured hemodynamic signal (Friston et al., 1994; Holmes and Friston, 1998). This allows to infer significant effects of cognitive states on brain activation, based on certain assumptions about the measured signal. As different GLMs can lead to different conclusions regarding experimental effects (Andrade et al., 1999; Carp, 2012), proper model assessment and model comparison is critical for statistically valid fMRI data analysis (Razavi et al., 2003; Monti, 2011).

In previous work, we have proposed *cross-validated Bayesian model selection* (cvBMS) to identify voxel-wise optimal models at the group level and then restrict group-level analysis to the best model in each voxel which avoids underfitting and overfitting in GLM-based fMRI data analysis (Soch et al., 2016). This approach allows for traditional analysis of neuroimaging data, but uses Bayesian inference for methodological control of such classical analyses. Importantly, cvBMS detects the model which is optimal in the majority of subjects, but not necessarily in all of them. Therefore, while this approach is powerful in a lot of cases and optimal in a decision-theoretic sense, it is not necessary in other cases and more appropriate alternatives exist.

These cases could, for example, differ by whether *regressors of interest* are contained in all models to be compared. By “regressors of interest”, we refer to those predictors whose estimates enter second-level analysis after first-level estimation. If these regressors are not part of all models in the model space, e.g. because the models differ by a categorical vs. parametric description of the experiment using completely different predictors (Bogler et al., 2013), one must employ the same model in all subjects in order to perform a sensible group analysis. In this case, cvBMS is the method of choice.

However, if regressors of interest are contained in all models, e.g. because the models only differ by *nuisance regressors* describing processes of no interest (Meyer and Haynes, in prep.), each model provides estimates for the parameters going into group analysis. In this case, cvBMS might unnecessarily lead one to use a model that is optimal in most subjects, but still sub-optimal in a lot of them. One could therefore speculate about performing second-level analysis on parameter estimates from different first-level models, depending on which model is optimal in each subject and voxel, which could be easily implemented using subject-wise selected-model maps (Soch et al., 2016).

In the present work, we generalize this idea to motivate a model averaging approach, more precisely a form of *Bayesian model averaging* (BMA). In fMRI data analysis, BMA has been described for dynamic causal models (Penny et al., 2010), but not so far for general linear models (Penny et al., 2007). In BMA, estimates of the *same* parameter from *different* models are combined with the models’ posterior probabilities (PP) and give rise to *averaged* parameter values which are more precise than *individual* models’ estimates. These averaged first-level parameters then enter classical second-level analyses. In our implementation, we calculate PPs from *cross-validated log model evidences* (cvLME) and refer to this as *cross-validated Bayesian model averaging* (cvBMA).

The rest of this paper falls into three parts. In Section 2, we describe the mathematical details of cvBMA for GLMs in fMRI, resting on both classical and Bayesian inference for the GLM. In Section 3, we apply cvBMA to simulated data and show that it leads to parameter estimates which are closer to their true values than estimates from the group-level or subject-wise best model. In Section 4, we apply cvBMA to empirical data before we discuss our results. Again, we show that cvBMA can be more sensitive than just using even the best GLM in each subject. Moreover, we find that cvBMS can prevent not detecting established effects, e.g. when using only one model.

## 2 Theory

### 2.1 The general linear model

As linear models, GLMs for fMRI (Friston et al., 1994; Kiebel and Holmes, 2011) assume an additive relationship between experimental conditions and the fMRI BOLD signal, i.e. a linear summation of expected hemodynamic responses into the measured hemodynamic signal. Consequently, in the GLM, a single voxel’s fMRI data (*y*) are modelled as a linear combination (*β*) of experimental factors and potential confounds (*X*), where errors (*ε*) are assumed to be normally distributed around zero and to have a known covariance structure (*V*), but unknown variance factor (*σ*^2^):

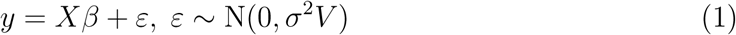

In this equation, *X* is an *n* × *p* matrix called the “design matrix” and *V* is an *n* × *n* matrix called a “correlation matrix” where *n* is the number of data points and *p* ist the number of regressors. In standard analysis packages like Statistical Parametric Mapping (SPM) (Ashburner et al., 2016), *V* is typically estimated from the signal’s temporal auto-correlations across all voxels using a Restricted Maximum Likelihood (ReML) approach (Friston et al., 2002b, a). In contrast to that, *X* has to be set by the user. Especially if regressors are correlated with each other, there can be doubt about which model to use. The general linear model (1) implicitly defines the following likelihood function:

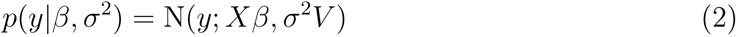

GLMs are typically inverted by applying maximum likelihood (ML) estimation to equation (2). This leads to ordinary least squares (OLS) estimates (Bishop, 2007, eq. 3.15) if *V* = *I_n_*, i.e. under temporal independence, or weighted least squares (WLS) estimates (Koch, 2007, eq. 4.29) if *V* ≠ *I_n_*, i.e. when errors *ε* are not assumed independent and identically distributed (i.i.d.).

Based on these ML estimates, statistical tests can be performed to investigate brain activity during different experimental conditions. These tests however strongly depend on the design matrix of the underlying model (Carp, 2012). When events overlap in time or are closely spaced temporally, convolution with the hemodynamic response function (Henson et al., 2001) will lead to positive correlation between the corresponding regressors. This influences parameter estimates for regressors of interest which in turn influences statistical tests and can change non-significant to significant or vice versa.

For mathematical convenience, we will rewrite the likelihood function (2) as

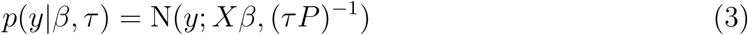

In this equation, *P* = *V*^−1^ is a *n* × *n* precision matrix and *τ* = 1/*σ*^2^ is the inverse residual variance (Koch, 2007, eq. 4.116). For Bayesian inference, it is advantageous to use the conjugate prior relative to equation (3). We have described this model, the general linear model with normal-gamma priors (GLM-NG), earlier and derived posterior distributions on the model parameters (Soch et al., 2016, eqs. 6) as well as the log model evidence for model comparison (Soch et al., 2016, eqs. 9).

In the following, we will introduce this model quality criterion (Section 2.2) and show how it can give rise to averaged model parameters (Section 2.3).

### 2.2 The log model evidence

Consider Bayesian inference on data *y* using model *m* with parameters *θ*. In this case, Bayes’ theorem is a statement about the posterior density (Gelman et al., 2013, eq. 1.1):

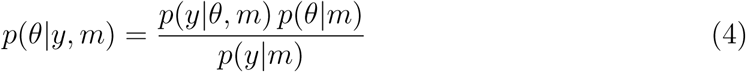

Here, *p*(*y|θ, m*) is the likelihood function, *p*(*θ|m*) is the prior distribution and the posterior distribution *p*(*θ|y, m*) is given as the normalized product of likelihood and prior. The denominator *p*(*y|m*) on the right-hand side acts as a normalization constant on the posterior density *p*(*θ|y, m*) and is given by (Gelman et al., 2013, eq. 1.3)

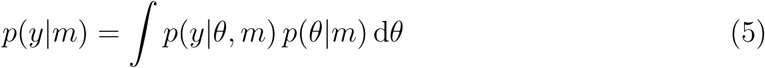

This is the probability of the data given only the model, independent of any particular parameter values. It is also called “marginal likelihood” or “model evidence” and can act as a model quality criterion in Bayesian inference (Penny, 2012). For computational reasons, only the logarithmized or log model evidence (LME) L(*m*) = log *p*(*y|m*) is of interest in most cases. For the GLM-NG, we have derived the posterior distribution (Soch et al., 2016, eq. 6) and log model evidence (Soch et al., 2016, eq. 9) in earlier work on model selection for GLMs.

The LME is a reliable model selection criterion as it (i) automatically penalizes for additional model parameters by integrating them out of the likelihood (Penny, 2012), (ii) can be naturally decomposed into model accuracy and model complexity (Penny et al., 2007) and (iii) accounts for the whole uncertainty about parameter estimates (Gelman et al., 2013) instead of using point estimates like classical information criteria such as AIC (Akaike, 1974) and BIC (Schwarz, 1978).

The LME however also requires prior distributions on the model parameters and typically diverges with ML-style flat priors. As multi-session fMRI data provides a natural basis for cross-validation (CV), we therefore suggested to use the LME in conjunction with CV (Soch et al., 2015) in order to avoid the necessity to specify prior distributions (see Figure S1) which are usually hard to come up with in fMRI research.

In this procedure, a posterior distribution is estimated from all sessions *j* ≠ *i* using non-informative prior distributions and then used as an informative prior distribution on the remaining session *i* to calculate the LME for this session (Soch et al., 2016). This is repeated for all CV folds, i.e. for each left-out session *i*, and the sum of out-of-sample LMEs (oosLME) then gives rise to the cross-validated LME (cvLME):

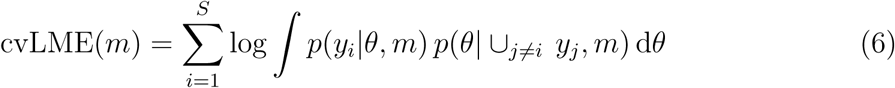

The cvLME has been validated using extensive simulations (Soch et al., 2016) and is insensitive to the number of folds into which the data are partitioned for cross-validation (see Figure S2; Soch and Allefeld, in prep.).

### 2.3 Bayesian model averaging

As log model evidences (LME) represent conditional probabilities, they can be used to calculate posterior probabilities (PP) using Bayes’ theorem (Hoeting et al., 1999, eq. 2):

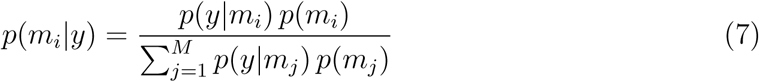

Here, *M* is the number of models, *p*(*y|m_i_*) is the *i*-th model evidence where the exponentiated cvLME has to be plugged in and *p*(*m_i_*) is the *i*-th prior probability which is usually set to *p*(*m_i_*) = 1/*M* making all models equally likely *a priori.* In this latter case, posterior probabilities are obtained as normalized exponentiated LMEs. Conceptually, this approach uses the first-level model evidences as the second-level likelihood function to make probabilistic statements about the model space.

After model assessment using LMEs, the PPs can be used to calculate averaged parameter estimates by performing Bayesian model averaging (BMA) (Hoeting et al., 1999, eq. 1):

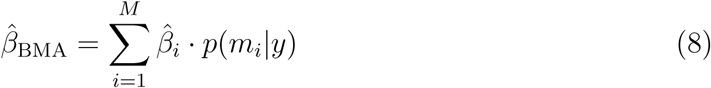

Here, 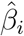 is the *i*-th model’s parameter estimate for a specific regressor and *p*(*m_i_|y*) is the *i*-th model’s posterior probability. Formally, BMA estimates can be seen as the parameter estimates of a larger model in which the variable “model” has been marginalized out. Note that, when one model is highly favored by the LME with a PP close to one, BMA is equivalent to just selecting this model’s parameter estimate. However, BMA automatically generalizes to cases where LMEs are less clear.^1^

BMA is theoretically advantageous (Hoeting et al., 1999) by making use of the whole posterior distribution across models, thereby accounting for modelling uncertainty, and has been empirically shown (Raftery et al., 1997) to provide improved predictive performance, e.g. measured using a logarithmic scoring rule (Good, 1952).

Model averaging is especially useful in, but not restricted to cases when the regressor or regressors for which the averaged parameter values are calculated are contained in all models of the model space. For this reason, BMA is particularly interesting when having identical regressors of interest, but varying regressors of no interest potentially correlated to the regressors of interest, which is often the case in fMRI data analysis due to different possible nuisance variables (see Figure 1A). We will investigate such cases in simulation settings (see Section 3) as well as with empirical data (see Section 4).

**Figure 1.**
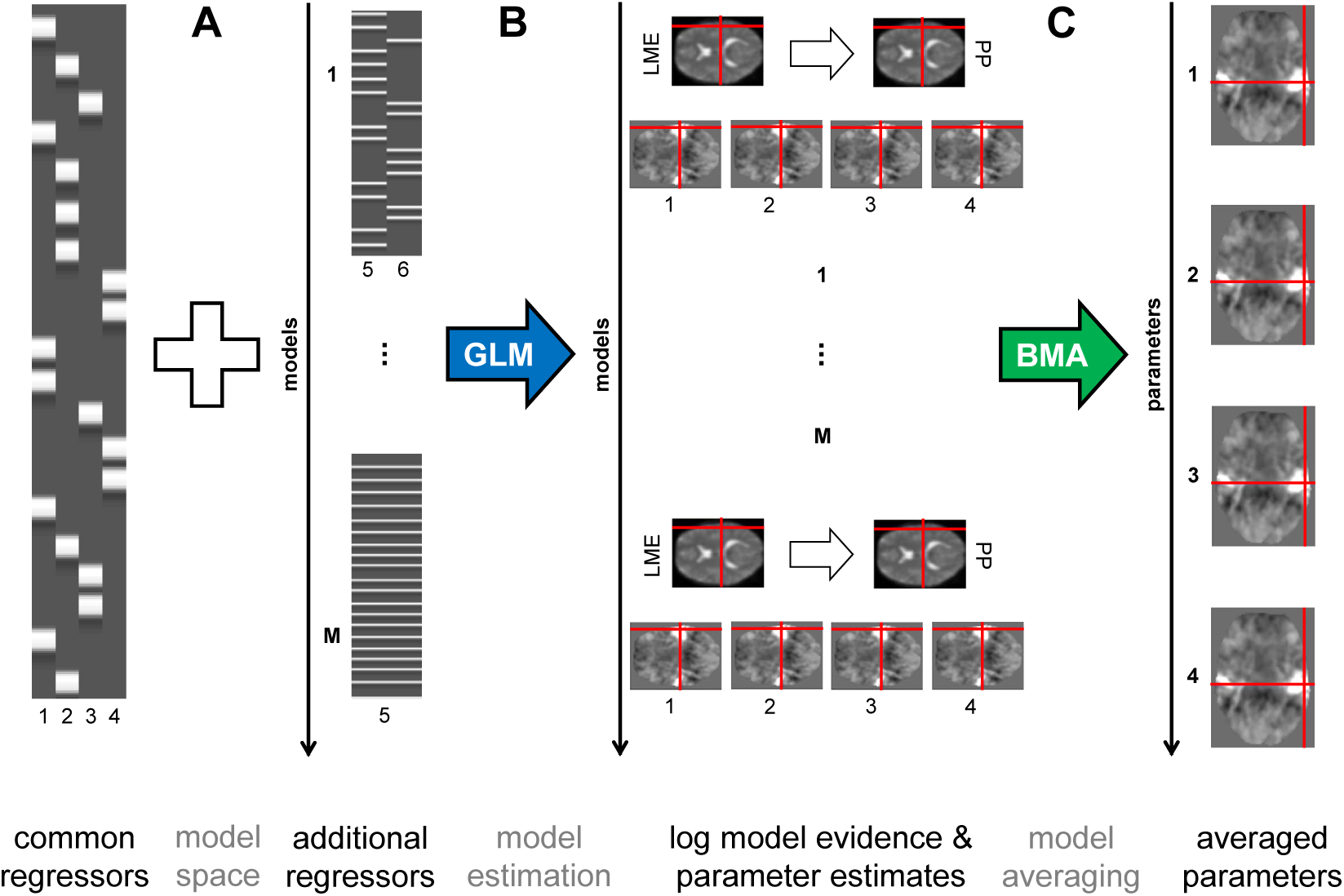
Model averaging for general linear models in fMRI data analysis. This figure summarizes our approach of cross-validated Bayesian model averaging (cvBMA). All calculations are performed voxel-wise, an exemplary voxel is highlighted using red crosshairs. (A) A model space is constructed by adding varying additional regressors (e.g. describing cues and feedback) to a set of common regressors (e.g. describing targets) which are included in all models. (B) Model estimation proceeds by classical CLM inversion, resulting in maximum likelihood (ML) parameter estimates (1-4), and by Bayesian CLM inversion with cross-validation (CV), resulting in maps of cross-validated log model evidences (LME) from which posterior probabilities (PP) can be calculated. (C) Model averaging proceeds by weighting different models’ estimates for the same regressor with the corresponding PP to obtain BMA estimates on which second-level inference can be performed. Parts of this figure are adapted from SPM course material (Stephan, 2010).

For the application of BMA to GLMs for fMRI, we use voxel-wise maximum-likelihood parameter estimates from SPM’s first-level analysis (Ashburner et al., 2016) and voxelwise cross-validated log model evidences as described earlier (Soch et al., 2016) (see Figure 1B). As we show in the Appendix, maximum likelihood (ML) estimates are equivalent to maximum a posteriori (MAP) estimates when non-informative priors are used. Finally, voxel-wise BMA estimates are calculated according to equation (8) (see Figure 1C). Together, this is referred to as cross-validated Bayesian model averaging (cvBMA).

## 3 Simulation

### 3.1 Methods

We test the cvBMA approach using simulated data and specifically investigate the impact of regressor correlation on various parameter estimation methods.

To this end, we imagine three different regressors: a “target” regressor specifying event onsets for a condition of interest (*x*_1_), a “cue” regressor with event onsets before targets (*x*_2_) and a “feedback” regressor with event onsets after targets (*x*_3_). Importantly, these experimental events have close temporal proximity so that convolution with the hemodynamic response function (HRF) leads to non-orthogonality. Here, we use a trial duration (*t*_dur_) of 2 s and modulate the onset difference (Δ*t*) from 6 s down to 2 s.

Based on these regressors, we define four different models: one consisting of only the target regressor (*m*_1_), two having the target regressor with either the cue regressor (*m*_2_) or the feedback regressor (*m*_3_) and one containing all three regressors (*m*_4_). In the full model *m*_4_, this leads to covariation of targets *x*_1_ with cues *x*_2_ and feedbacks *x*_3_ where correlation between regressors decreases with onset difference (see Figure 2A).

**Figure 2.**
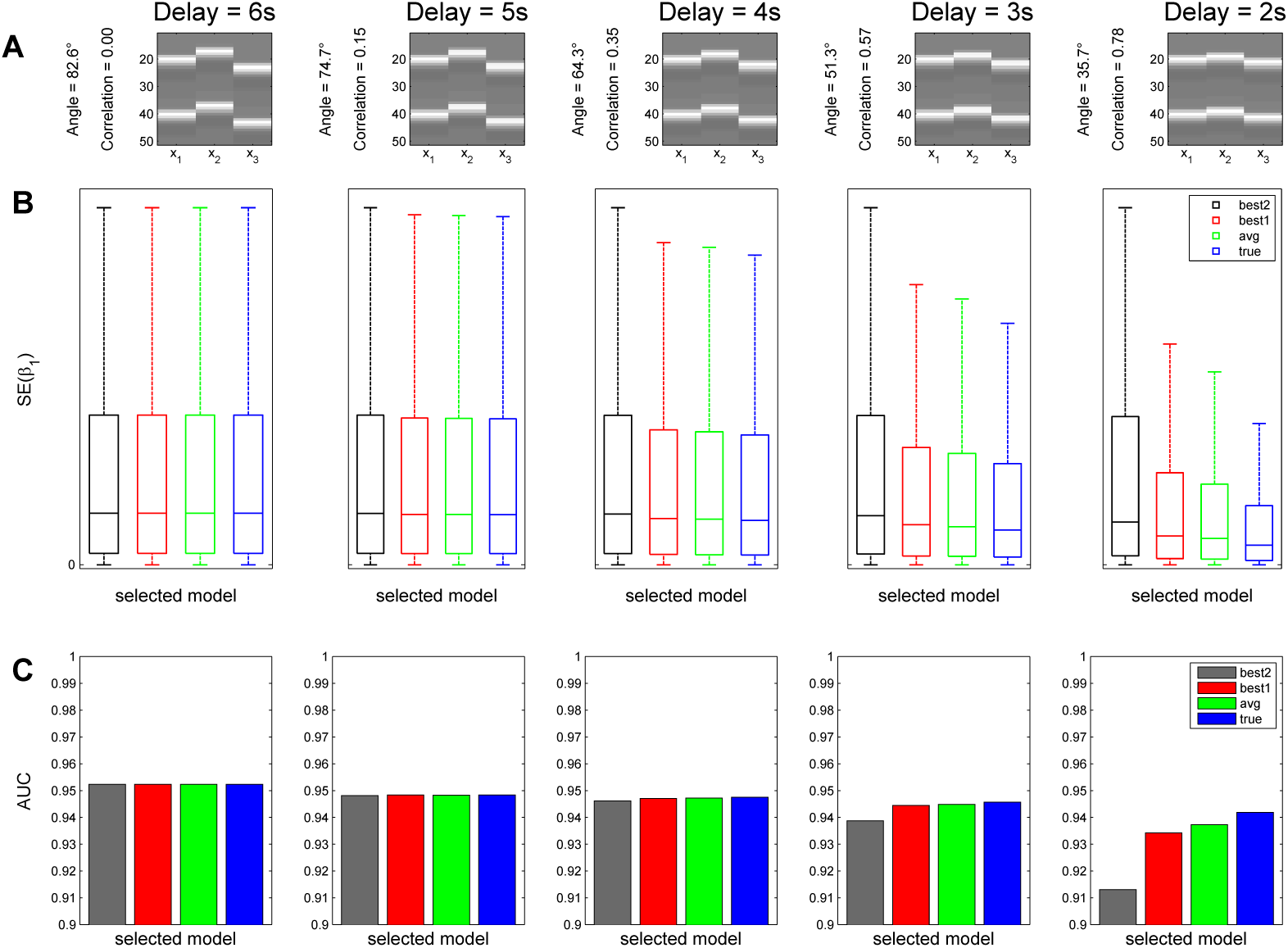
Simulation performance of cross-validated Bayesian model averaging. This figure demonstrates that Bayesian model averaging (BMA) can be superior to always using parameters from the best model as identified by maximal log model evidence (LME). (A) Design matrices of the full model used in the simulation. Different onset differences between a target regressor (*x*_1_) and preceding cues (*x*_2_) as well as subsequent feedback (*x*_3_) are simulated where a delay in seconds implies a certain angle in degrees and correlation between *x*_1_ and *x*_2_ as well as *x*_1_ and *x*_3_. (B) Box plots of squared errors (SE) across simulations when estimating the target regressor weight (*β*_1_) using either the group-level best model (obtained by cvBMS, black), the subject-wise best model (with maximal cvLME, red), the averaged model (obtained by cvBMA, green) or the true model (used to generate the data, blue). Each panel is scaled such that the upper black whisker corresponds to 95% of the y-axis maximum and the y-axis minimum is at zero. (C) Bar plots of area under the curve (AUC) when performing an ROC analysis for the second-level one-sample t-test of *β*_1_ against 0. Each panel is scaled to 0.9 *<* AUC *<* 1. When regressors are almost orthogonal, the four estimation techniques do not differ regarding SE or AUC. With decreasing delay and thus increasing overlap, the true model outperforms cvBMA and cvBMA outperforms the best-model approaches in terms of both, estimation precision (B) and test performance (C).

For each onset difference or delay, *N*_1_ = 10,000 samples with *N*_2_ = 25 subjects per sample are simulated as follows. *First*, a true model is randomly drawn from *M* = {*m*_1_, *m*_2_, *m*_3_, *m*_4_} for each subject. For each model and delay, design matrices *X* for *S* = 5 sessions were generated before simulation. Each session consisted of *n* = 200 scans at TR = 2 s containing 9 trials with a duration of 2 s in intervals of 20 scans and the respective delay between target and cue or feedback (see Figure 2A).

*Second*, true regression coefficients are drawn using the relation

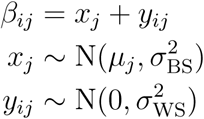

where *i* and *j* index session and parameter respectively, *μ_j_* is the true population average of one regressor’s effect, 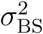 represents the subject-to-subject variance and 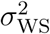 represents the session-to-session variance.

Following a bottom-up approach, the scan-to-scan variance *σ*^2^ was set to 1. Then, the ratio of between-subject variance to within-subject variance 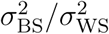 was set to 2 (cf. Soch et al., 2016, fig. 6C) and the variances were chosen such that the full model *m*_4_ exhibited an expected signal-to-noise ratio 〈SNR〉 = 〈var(*Xβ*)/*σ*^2^〉 of 0.1. This resulted in values 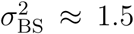 and 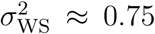. Finally, the population mean *μ*_1_ for the target effect *β*_1_ was calibrated such that the power of a one-sample t-test against *μ*_0_ = 0 across *N*_2_ = 25 subjects drawn from a population with total variance of 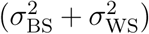 was 0.8 at a significance level of *α* = 0.05. This resulted in the value *μ*_1_ ≈ 0.75. For the confound effects *β*_2_ and *β*_3_, *μ* was set to 0 reflecting that confound effects exist, but that they are unsystematic across subjects.^2^

With *μ*_1_ ≠ 0, simulations allows to assess the true positive rate (TPR), i.e. statistical power, of a one-sample t-test of *β*_1_ against 0. In another *N*_1_ = 10,000 simulations, *μ*_1_ was simply set to 0 which allows to assess the false positive rate (FPR), i.e. type I error probability, of the same statistical test.

*Third*, simulated data are generated by sampling zero-mean Gaussian observation noise *ε* ~ N(0, *σ*^2^*V*) where the temporal covariance *V* was set to invoke fMRI-typical autocorrelations^3^ and then adding the random noise to the true signal to get a measured signal *y* = *Xβ* + *ε* that was entered into analyses.

Finally, as the target regressor (*x*_1_) was included in all design matrices of our model space, the model parameter corresponding to target presentation (*β*_1_) was estimated using all four models and models were quantified using the cross-validated log model evidence (cvLME). Then, we compared four parameter estimates for *β*_1_: the one obtained using the group-level best model (obtained by cvBMS), using the subject-wise best model (with maximal cvLME), using Bayesian model averaging (obtained by cvBMA) and using the true model (used to generate the data).

### 3.2 Results

The impact of covariation between regressors on parameter estimates in the general linear model (GLM) using ordinary least squares (OLS) is best captured using the inner product of these regressors. The normalized inner product of two vectors is equal to the cosine of the angle between them. With 6 s delay, regressors are almost orthogonal, i.e. their angle is close to 90°. With a delay of 2 s, we observe an angle of 35.7° and a correlation of 0.78 in our simulations (see Figure 2A), implying a considerable degree of non-orthogonality, but not collinearity between target and cue or feedback regressors.

The precision of parameter estimates can be described by squared errors (SE), i.e. squared differences between true and estimated parameter values. We were interested in SE(*β*_1_), because the parameter estimate for the target regressor could possibly be confounded by the cue and feedback regressors, depending on whether they were part of the true model or not. Additionally, the TPR for a t-test of *H*_1_:*β*_1_ > 0 against *H*_0_: *β*_1_ = 0 was plotted against the FPR in a receiver-operating characteristic (ROC) analysis to calculate the area under the curve (AUC) as a measure of statistical test performance.

First, in the case of a large onset difference, the SEs for this parameter are the same for all models, because additional regressors do not change parameter estimates if they are orthogonal to the regressor of interest (see Figure 2B, left panels). Second, when there is temporal overlap between event regressors, SEs are on average smallest when the true model is used (see Figure 2B, blue bars), because the model used to generate the data leads to the most precise parameter estimates. Third, while the true model slightly outperforms cvBMA estimates, cvBMA estimates slightly outperform the subject-wise best model and strongly outperform the group-level best model (see Figure 2B, green/red/black), because BMA accounts for the whole uncertainty over models and does not just select from the model with maximal LME. Interestingly, the best model and the averaged model get very close to the true model in cases of moderate correlation which suggests that LMEs are very decisive so that the advantage of BMA is not very high. This is also reflected in AUC values not differing very much between models for medium overlap (see Figure 2C).

The results demonstrate that Bayesian model averaging, i.e. weighting parameter estimates according to the models’ posterior probabilities, can be better than using the best individual model, i.e. taking parameter estimates from the model with maximal posterior probability. Although cvBMA is worse than using the true model, it is the optimal approach for empirical data, because the true model is unknown in such cases. These simulations are therefore a first indication for employing cvBMA in fMRI data analysis. A second indication will be provided by the analysis of empirical data.

## 4 Application

### 4.1 Methods

We test the cvBMA approach using empirical data from a conflict adaptation paradigm (Meyer and Haynes, in prep.) and specifically investigate the capability of model averaging to identify experimental effects that would be undetectable by individual models.

The experimental paradigm (see Figure 3) was an Eriksen flanker task (Eriksen and Eriksen, 1974) combined with a response rule switch (Bode and Haynes, 2009) giving a 2 x 2 factorial design, the two factors being conflict (congruent vs. incongruent) and task set (response rule 1 vs. 2). In each trial, three vertical arrows (pointing upward or downward) were presented at the center of the screen. The upper and the lower arrow were either pointing in the same (congruent) or in opposite direction (incongruent) when compared to the target arrow in the center (see Figure 3C). Subjects were requested to indicate the direction of the target arrow via right-hand button press, but the response rule was changing from block to block (see Figure 3C).

**Figure 3.**
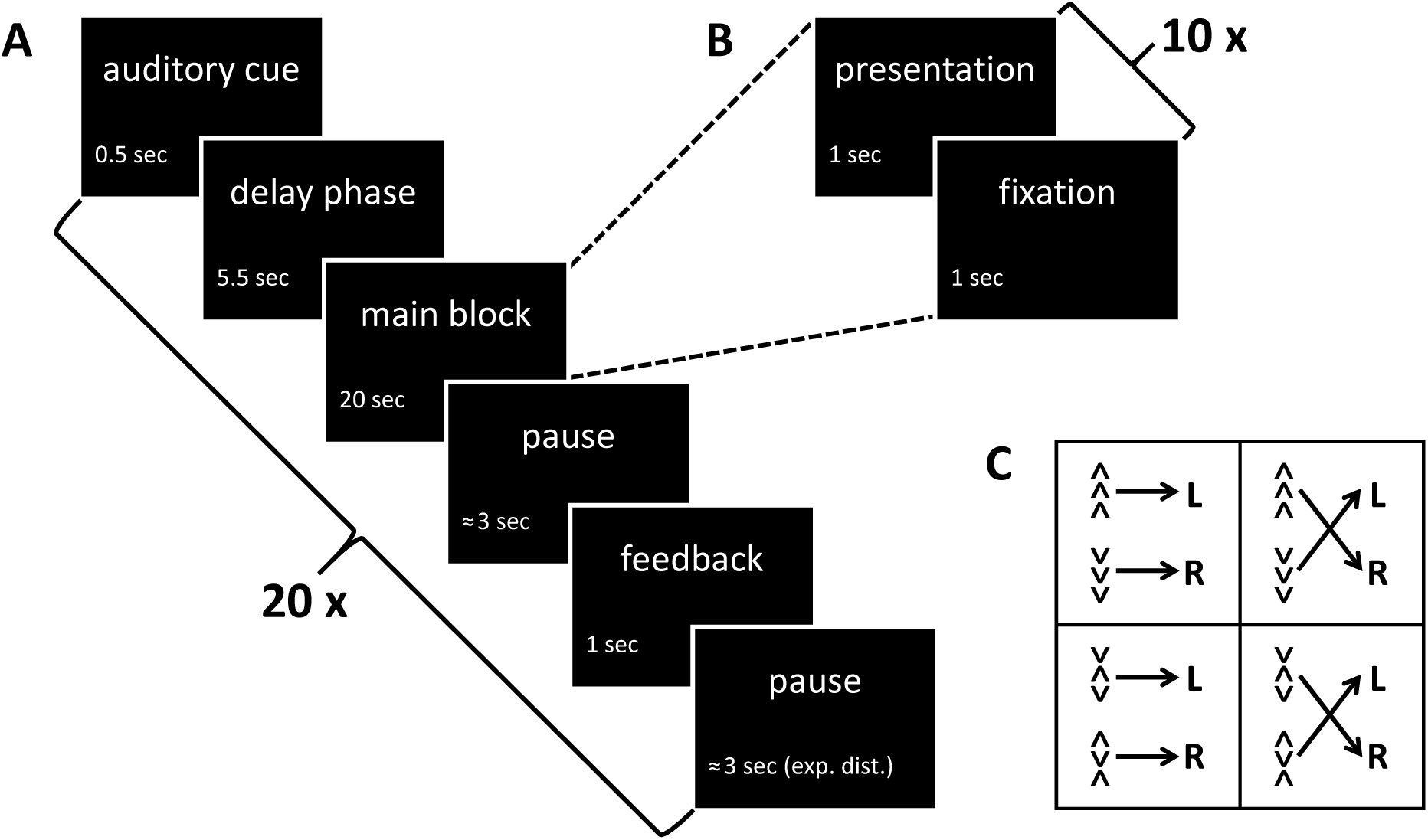
Experimental design of the conflict adaptation paradigm. This figure describes the experiment underlying the data set used for empirical validation of our method. (A) Sequence of events and exact timing during each of the 20 blocks per session. (B) Sequence of events and exact timing during one of the 10 trials per block. (C) Experimental conditions: The paradigm was a 2 × 2 design with conflict (congruent vs. incongruent) and task set (response rule 1 vs. 2) being the two factors. Abbreviations: sec = seconds; exp. dist. = exponentially distributed; L/R = left/right button.

Stimuli were presented in 6 sessions with 20 blocks of 10 trials. Each trial consisted of 1 second presentation and 1 second fixation (see Figure 3B). Each block lasted 10 x 2 = 20 seconds, was preceded by a 500 ms auditory cue signalling the response rule as well as a 5.5 second delay phase to prepare for the coming block and was succeeded by a 1 second feedback phase indicating how many blocks would still follow (see Figure 3A). There were 5 blocks per each of the 4 conditions. The total duration of one block was around 33 seconds and each fMRI session was lasting 666 seconds.

fMRI data were preprocessed using SPM12, Revision 6225 per 01/10/2014 (http://www.fil.ion.ucl.ac.uk/spm/software/spm12/). Functional MRI scans were corrected for acquisition time delay (slice timing) and head motion (spatial realignment), normalized to MNI space using a voxel size of 3 x 3 x 3 mm (spatial normalization), smoothed using a Gaussian kernel with a full width at half maximum (FWHM) of 6 x 6 x 6 mm (spatial smoothing) and filtered using a high-pass filter (HPF) with cut-off at *T* = 128 s (temporal filtering). Unless otherwise stated, SPM12 default parameters were used.

For first-level analysis, we categorized each block as containing congruent or incongruent stimuli and by whether the response rule switched or stayed relative to the preceding block.^4^ This lead to four categories of blocks: congruent-stay, congruent-switch, incongruent-stay, incongruent-switch. For each subject and each session, a GLM was specified including four regressors modelling these four types of blocks and two regressors modelling delay phases for switch blocks and for stay blocks. There was no modelling of error trials, button presses, reaction times or feedback phases. Each model included six movement parameters obtained from spatial realignment. A first-order auto-regressive AR(1) model was used to account for noise auto-correlations.

In second-level analysis, we were interested in the different neural activity in the switch-delays preceding the application of a new response rule and the stay-delays preceding the application of the same response rule as before. To again induce correlation between regressors, we introduce two variable model space features with regressors overlapping with the delay phase regressors and therefore influencing their estimates.

First, the 500 ms auditory cues before the 5.5 second delay phases were modelled by two extra regressors, also separated by the switch-stay difference. This model feature was motivated by the fact these stimulations indeed require two different cognitive processes, namely auditory perception for the cues and executive planning for the delays.

Second, the first trials of stimulation blocks were additionally modelled by four extra regressors, separated exactly like the block regressors. This model feature was motivated by the fact that the first trial of each block could demand a restart cost which was also observable in the distribution of reaction times, i.e. a significantly higher reaction time in the first trial compared to later trials (Meyer and Haynes, in prep.).

Taken together, this resulted in four possible models for first-level data (see Table 2): all of them with blocks and delays being modelled; one with only cue phases being additionally modelled, one with only first trials being additionally modelled, one with both and one without both. Across all subjects and sessions, average correlation between delays and cue phases was 0.63 and average correlation between delays and first trials was 0.46. Like in our simulation study, the goal was to investigate the properties of statistical inference being performed with individual models, using the best GLM as identified by maximal cvLME and using BMA estimates based on the models’ cvLMEs.

### 4.2 Results

On the second level, we focused on the delay phases and looked for a main effect of stay vs. switch blocks. We hypothesized that delay phases after cues indicating a switch of the response rule might elicit preparatory processes that lead to higher motor cortex activity in the left hemisphere (participants responded with their right hand) compared to delay phases preceding blocks with the same response rule as before. Such an effect has been suggested by early work on task preparation (Brass and Cramon, 2002), experimentally demonstrated (Kim et al., 2011) and meta-analytically validated (Kim et al., 2012). In fact, this main effect later turned out to be due to a positive effect of switch over stay blocks (see Table 1, right-hand side).

**Table 1.**
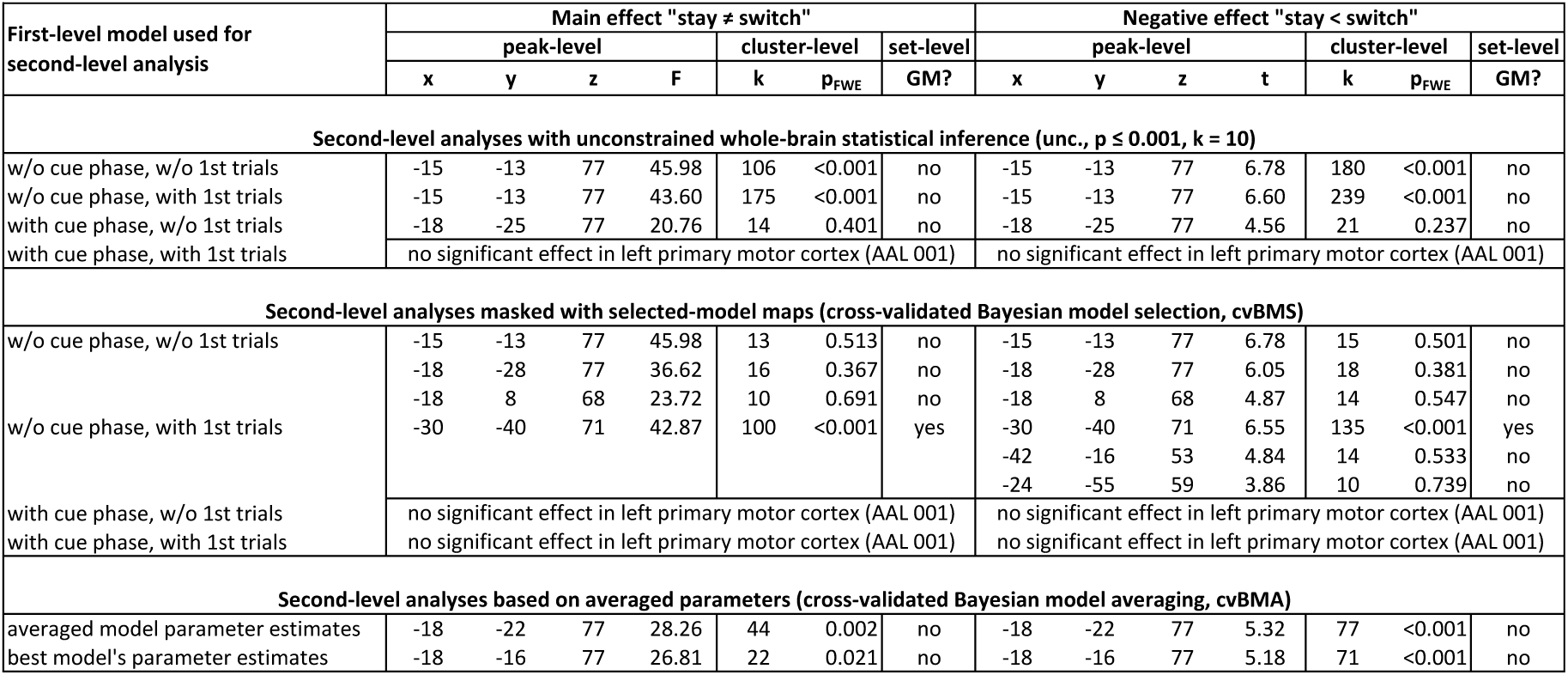
Empirical example for cross-validated Bayesian model averaging. In each row, peak-level, cluster-level and set-level statistics are given for an F-test of the main effect between stay and switch blocks as well as a t-test of the negative effect of stay against switch blocks. The upper section of the table summarizes unconstrained whole-brain statistical inference using the four models. The effect in question can be detected using three models, but not the most complex one. The middle section of the table summarizes second-level analyses that were masked using selected-model maps (SMM) from cross-validated Bayesian model selection (cvBMS). The effect in question can be detected using the two models that do not include cue phase regressors. The lower section of the table summarizes second-level analysis based on averaged parameters (cvBMA estimates) and the subject-wise best GLM’s parameter estimates (maximal cvLME). The effect in question can be detected using both methods, but is stronger when employing the cvBMA approach. Abbreviations: *x, y, z* = MNI coordinates; *F/t* = F-/t-statistic, *k* = cluster size; pFWE = family-wise error-corrected p-value; GM = global maximum.

First, we tried to identify this effect using the four first-level models as such. We performed second-level analysis using the summary-statistic approach (Holmes and Friston, 1998) and were able to detect a main effect of stay vs. switch in left primary motor cortex using all models except the one modelling both, cue phases and first trials (see Table 1, upper section). The effect was not significant at the cluster level under correction for family-wise errors (FWE) when using the model with cue phases but without first trials, indicating that modelling the cue phase had a higher impact on parameter estimates for the delay phase due to their shared variance and higher correlation, making the difference between stay and switch blocks insignificant.

Next, we performed cross-validated Bayesian model selection (cvBMS) to identify the group-level optimal model in each voxel (Soch et al., 2015) and observed that all models are optimal in at least some voxels of the left precentral gyrus (see Table 2). We used this information to generate selected-model maps (SMM) indicating for each model in which voxels it is optimal and masked second-level analyses using these SMMs in order to restrict statistical inferences to those voxels where the corresponding model is best explaining the data at the group level. This approach, as suggested in previous work (Soch et al., 2016), lead to the effect only being detected by the models not accounting for cue phases, again suggesting that modelling these had the greater influence on delay phase significance (see Table 1, middle section). In the model including first trials but not cue phases, the main effect of stay vs. switch blocks in left primary motor cortex was also the global maximum on the respective contrast. This demonstrates that cvBMS can prevent us from overfitting and not detecting established effects when using just one model. Notably, the most complex model including both, cue phases and first trials, would not have been the best choice here.

**Table 2.**
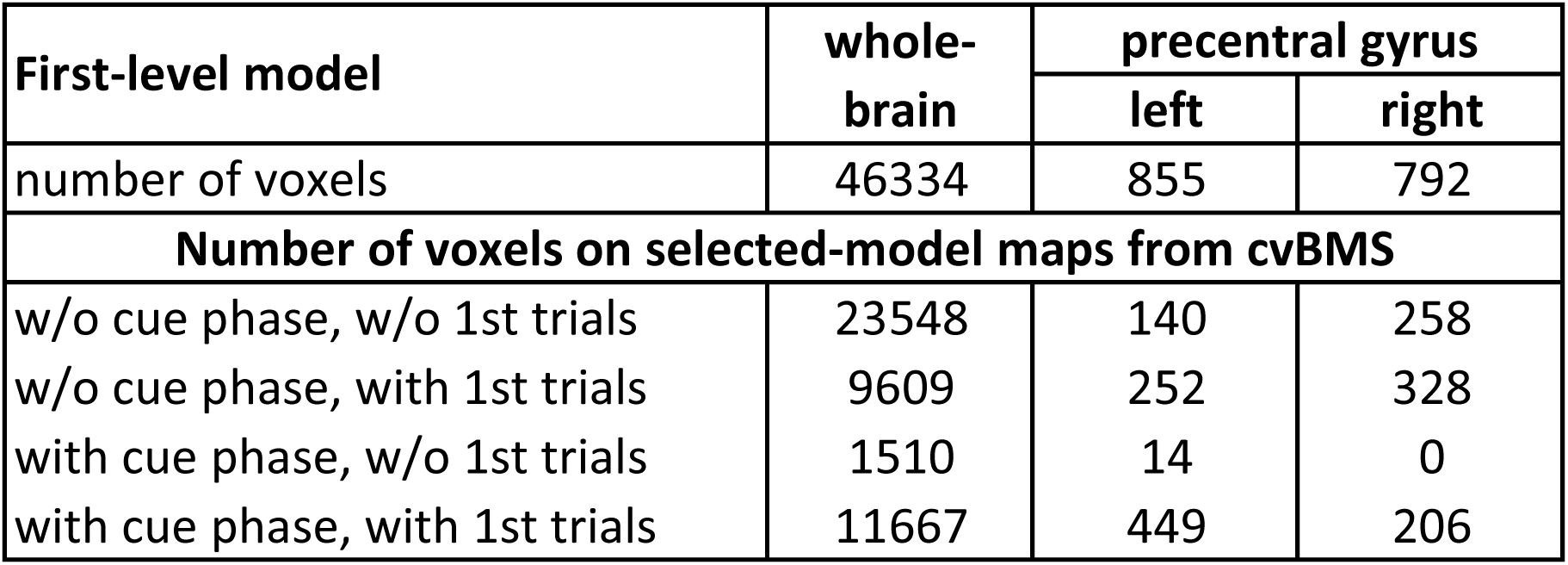
Selected models from cross-validated Bayesian model selection. This table summarizes model selection results for the four models used in the empirical example (see Figures 3 and 4). For each model, the number of voxels in which it is selected as the optimal model by cross-validated Bayesian model selection (cvBMS) is given for the left and right precentral gyrus, taken from the Automated Anatomical Labeling (AAL) atlas (AAL 001 and AAL 002) and putatively including the primary motor cortices, as well as at the whole-brain level.

| First-level model | whole-brain | precentral gyrus |  |
| --- | --- | --- | --- |
|  |  | left | right |
| number of voxels | 46334 | 855 | 792 |
| **Number of voxels on selected-model maps from cvBMS** |  |  |  |
| w/o cue phase, w/o 1st trials | 23548 | 140 | 258 |
| w/o cue phase, with 1st trials | 9609 | 252 | 328 |
| with cue phase, w/o 1st trials | 1510 | 14 | 0 |
| with cue phase, with 1st trials | 11667 | 449 | 206 |

Last, we compared two estimation methods not being based on individual models: using parameter estimates from the subject-level optimal models as identified by maximal LME and using cross-validated Bayesian model averaging (cvBMA) as developed in the present work. For both methods, voxel-wise cvLMEs were calculated for each model in each subject. For the best-model approach, first-level parameter estimates in each voxel were taken from the model having the highest cvLME in this voxel and then subjected to second-level analysis. For the model averaging approach, first-level parameter estimates in each voxel were weighted according to posterior probabilities calculated from the models’ cvLMEs in this voxel (see Figure 1) and the averaged parameters were subjected to second-level analysis. The best-model approach is equivalent to setting the posterior probability of the most likely model to 1 and then performing Bayesian model averaging.

We observe that the effect in question can be identified using both methods, but with a different degree of significance. When test statistic, cluster size or p-value are taken as a measure of sensitivity, cvBMA must be judged superior to using the subject-wise best GLM (see Table 1, lower section). Interestingly, cvBMA does not only calculate the weighted average of the models’ parameter estimates per voxel, but also seems to make a compromise regarding the spatial location of the main effect when compared to statistical inferences based on the individual models (see Figure 4A).

**Figure 4.**
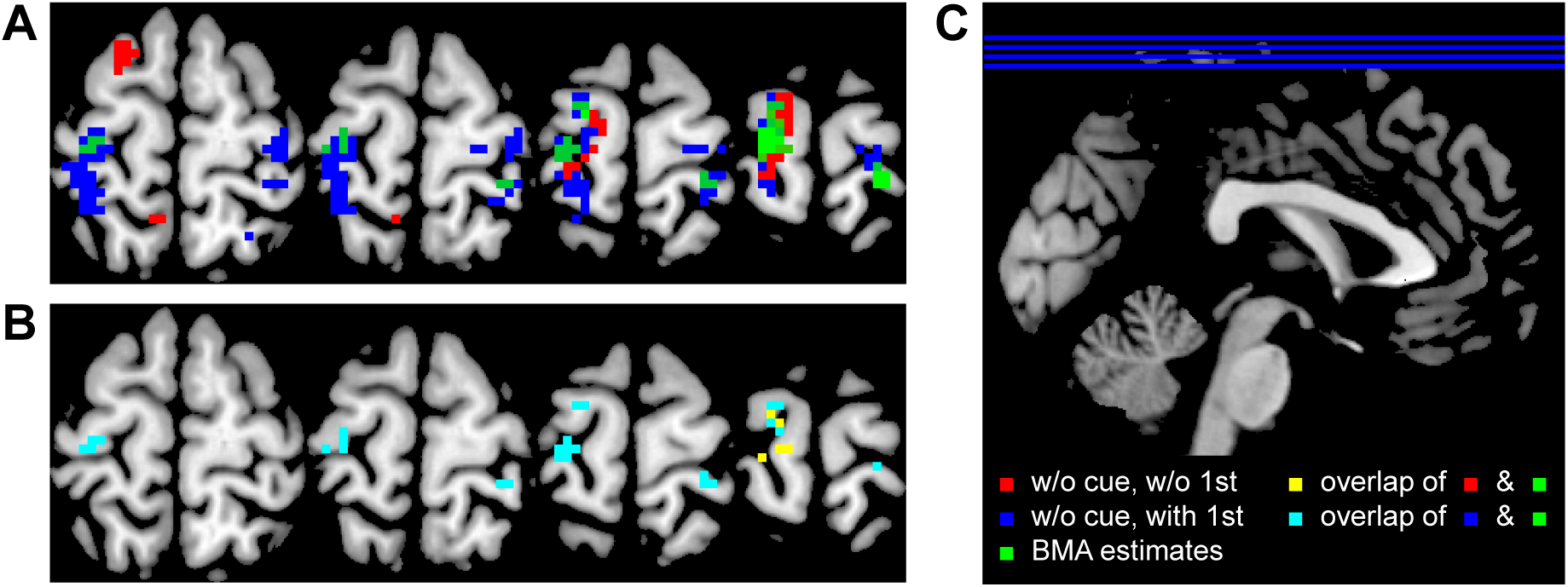
Empirical example for cross-validated Bayesian model averaging. This figure illustrates how model averaging achieves a compromise between the different models’ parameter estimates. (A) Colored voxels indicate a significant main effect of stay vs. switch blocks observed when using a model without cue phases and first trials (red), a model without cue phases but with first trials (blue), both masked with their group-level selected-model map (SMM) obtained through cross-validated Bayesian model selection (cvBMS), as well as observed in a second-level analysis on the averaged parameters (green) obatined via cross-validated Bayesian model averaging (cvBMA). The BMA approach seems to not only average parameter values, but to also make a compromise regarding spatial location with respect to the blue (more ventral-posterior) and red (more dorsal-anterior) clusters. (B) Colored voxels indicate overlap between significant effects using individual models and cvBMA estimates. The BMA approach shows stronger overlap with the model including first trials (cyan) than with the one not including them (yellow). By definition, there is no overlap between these two models as SMMs are mutually exclusive. (C) Axial sections and color coding used for the panels on the left.

Like our simulation study, this empirical example therefore indicates that performing cvBMA can be better than using parameter estimates from the subject-wise best GLMs. Using cvBMA, we can improve parameter estimates for regressors of interest by drawing information from a variety of models instead of just relying on one particular model. This is possible, because each model provides parameter estimates for these regressors and because the models differ in how well they are supported by the measured data, as quantified by their posterior probabilties. As demonstrated in simulation, by factoring in our uncertainty in this way, parameter estimates move closer to their true values which in turn increases the sensitivity for experimental effects.

## 5 Discussion

We have introduced a model averaging approach for optimizing parameter estimates when analyzing *functional magnetic resonance imaging* (fMRI) data using *general linear models* (GLMs). We have demonstrated that *cross-validated Bayesian model averaging* (cvBMA) serves its intended purpose and that it is useful in practice. As nuisance variables and correlated regressors are common topics in fMRI data analysis, usage of this technique reduces model misspecification and thereby enhances the methodological quality of functional neuroimaging studies (Friston, 2009).

Often, psychological paradigms combined with fMRI use trials or blocks with multiple phases (e.g. cue – delay – target – feedback; see Meyer and Haynes, in prep.), so that the basic model setup (the target regressors) is fixed, but there is uncertainty about which processes of no interest (cues, delays, feedback) should be included into the model (Andrade et al., 1999). Especially in, but not restricted to these cases of correlated regressors (Mumford et al., 2015), cvBMA has its greatest potential which is why our simulated data and the empirical examples were constructed like this.

Typically, if one is unsure about the optimal analysis approach in such a situation, just one model is estimated or, even worse, a lot of models are estimated and model selection is made by looking at significant effects (Soch et al., 2016). Here, model averaging provides a simple way to avoid such biases. It encourages multiple model estimation in order to avoid mismodelling, but calculates weighted parameter estimates by combining the models in order to avoid subjective model selection. These weighted parameters can then be used for second-level analyses within standard workflows, e.g. SPM.

Using simulated data, we were able to show that averaged model parameter estimates have a smaller mean squared error than even the best model’s parameter estimates. Using empirical data, we demonstrated the trivial fact that different GLMs can lead to the same effect being either significant or insignificant. Interestingly, we found that the most complex GLM is not always the best, speaking against the fMRI practitioner’s maxim that the design matrix should “embody all available knowledge about experimentally controlled factors and potential confounds” (Stephan, 2010) – though it should still be applied in the absence of any knowledge about model quality.

Although the most complex model was not optimal in this case, our previously suggested approach of *cross-validated Bayesian model selection* (cvBMS) and subsequent masking of second-level analyses with selected-model maps (SMM) was able to protect against not detecting an established experimental effect which additionally validates this technique (Soch et al., 2016). Moreover, *cross-validated Bayesian model averaging* (cvBMA) was found to be more sensitive to experimental effects than simply extracting parameter estimates from the best GLM in each subject which again highlights its applicability in situations of uncertainty about modelling processes of no interest.

However, we want to caution the reader that model averaging is not a general remedy against confound effects. In cases of low and medium correlation, cvBMA can be a useful tool to adjust parameter estimates of experimental regressors for the effects of confound regressors. But if two HRF regressors are correlated by say 0.99, and just one of them has an effect on the data, it is almost impossible to find out where this effect really comes from. Any model selection will either increase the false positive rate (when including an experimental regressor which does not have an effect) or decrease the true positive rate (when including a confound regressor that does not have an effect) of statistical tests for experimental effects. Even model averaging can only achieve a compromise between these two suboptima. Also with model selection methods at hand, one should still try to avoid confounds in experimental designs. And if confounds are unavoidable *within* subjects, they should at least not be consistent *across* subjects.

Moreover, one has to keep in mind that cvBMS and cvBMA do not only perform different statistical operations, but also have different interpretations. Whereas cvBMS aims at identifying which psychological model best describes the hemodynamic signal, cvBMA tries to optimize decisions with respect to certain model parameters, in this case by improving parameter estimates. For example, if we look at Figure 4A, red and blue voxels indicate that the respective models are optimal *and* the respective contrast is significant in these voxels. In contrast, green voxels indicate that the respective contrast is significant in these voxels, *given* that model uncertainty has been removed.

All in all, we therefore see cvBMA as a complement to the recently developed cvBMS. While cvBMS is the optimal approach when parameters of interest are not identical across the model space, e.g. because one part of the models uses a categorical and another part uses a parametric description of the paradigm (Bogler et al., 2013), cvBMA is the more appropriate analysis when regressors of interest are the same in all models (Meyer and Haynes, in prep.), such that their estimates can be averaged across models and taken to second-level analysis for sensible population inference.

## 6 Acknowledgements

This work was supported by the Bernstein Computational Neuroscience Program of the German Federal Ministry of Education and Research (BMBF grant 01GQ1001C), the Research Training Group “Sensory Computation in Neural Systems” (GRK 1589/1-2), the Collaborative Research Center “Volition and Cognitive Control: Mechanisms, Modulations, Dysfunctions” (SFB 940/1) and the German Research Foundation (DFG grants EXC 257 and KFO 247).

Joram Soch received a Humboldt Research Track Scholarship and receives an Elsa Neumann Scholarship from the State of Berlin. The authors have no conflict of interest, financial or otherwise, to declare.

## 7 Software Note

Implementations of voxel-wise cross-validated Bayesian model selection (https://github.com/JoramSoch/cvBMS) and cross-validated Bayesian model averaging (https://github.com/JoramSoch/cvBMA) compatible with SPM8 and SPM12 can be downloaded from the corresponding author’s GitHub profile (https://github.com/JoramSoch).

## 8 Appendix

Consider the general linear model (GLM) given by

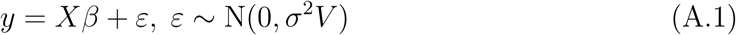

with known design matrix *X* and covariance structure *V* = *P*^−1^ as well as unknown model parameters *β* and *σ*^2^ = 1/*τ*. Then, maximum likelihood (ML) estimates for regression coefficients *β*, their covariance cov(β) and residual variance *σ*^2^ are given by

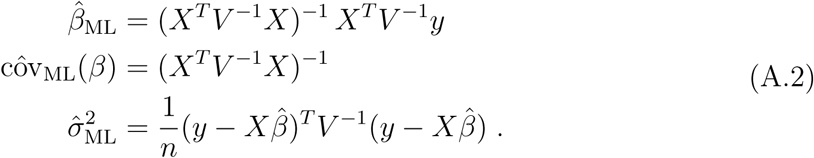

The most common Bayesian treatment of linear regression is the general linear model with normal-gamma priors (GLM-NG; Bishop, 2007, ch. 3.4; Koch, 2007, ch. 4.3.2). Using a non-informative prior distribution (Soch et al., 2016, eq. 15), the parameters of the posterior distribution (Soch et al., 2016, eq. 6) evaluate as

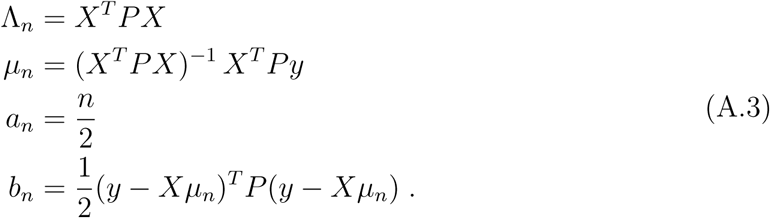

Then, maximum a posteriori (MAP) estimates for regression coefficients *β*, their covariance cov(*β*) and residual precision *τ* are given by

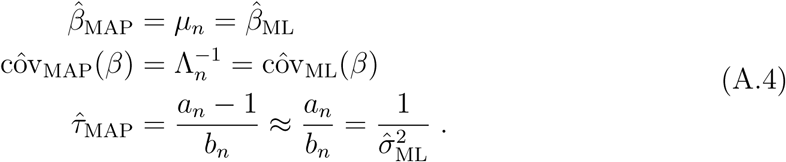

This demonstrates that ML estimates for the regression coefficients in one session correspond to MAP estimates when analyzing this session with non-informative priors. Given that all sessions contribute the same amount of evidence – which usually is the case in fMRI when sessions (approximately) use the same number of scans –, the MAP estimate from all data is also equal to the average of the session-wise ML estimates

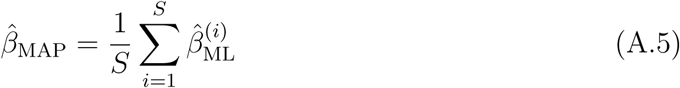

where *i* indexes session. It follows that BMA using averaged ML estimates (e.g. estimated using SPM) is equivalent to BMA using MAP estimates obtained with non-informative priors (e.g. when calculating the cvLME).

## Footnotes

1 Please note that the cvLME is used for model averaging and therefore session-wise parameter estimates (e.g. obtained from SPM) are averaged across sessions before applying the BMA equation. In the case of single-session fMRI data, we suggest to use split-half cross-validation (Soch et al., 2014) and use the cvLME together with the one estimate for each regressor.

2 For comparison, in an SPM template data set, we have observed the median values *μ* = 0. 14, 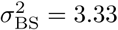 and SNR = 4.10, i.e. a much higher SNR than in our simulations. These data were first published as a study on repetition priming (Henson et al., 2002), previously used for model comparison (Penny et al., 2007) and analyzed according to the SPM8 Manual (Ashburner et al., 2013, ch. 29).

3 In our MATLAB script, *V* was created using the command: `V = toeplitz(exp(-[0:1: (n-1)]))`.

4 The first block was categorized as a switch block, because a new rule had to be applied.

